# Delta-1 functionalized hydrogel promotes hESC-cardiomyocyte graft proliferation and maintains heart function post-injury

**DOI:** 10.1101/768226

**Authors:** Kaytlyn A Gerbin, Katie A Mitzelfelt, Xuan Guan, Amy M Martinson, Charles E Murry

## Abstract

Current cell transplantation techniques are hindered by small graft size, requiring high cell doses to achieve therapeutic cardiac remuscularization. Enhancing the proliferation of transplanted human stem cell-derived cardiomyocytes (hESC-CMs) could address this, allowing an otherwise subtherapeutic cell dose to prevent disease progression after myocardial infarction. Here, we designed a hydrogel that activates Notch signaling through 3D presentation of the Notch ligand Delta-1 to use as an injectate for transplanting hESC-CMs into the infarcted rat myocardium. After four weeks, hESC-CM proliferation increased 2-fold and resulted in a 3-fold increase in graft size with the Delta-1 hydrogel compared to controls. To stringently test the effect of Notch-mediated graft expansion on long-term heart function, a normally subtherapeutic dose of hESC-CMs was implanted into the infarcted myocardium and cardiac function was evaluated by echocardiography. Transplantation of the Delta-1 hydrogel + hESC-CMs augmented heart function and was significantly higher at three months compared to controls. Graft size and hESC-CM proliferation were also increased at three months post-implantation. Collectively, these results demonstrate the therapeutic approach of a Delta-1 functionalized hydrogel to reduce the cell dose required to achieve functional benefit after myocardial infarction by enhancing hESC-CM graft size and proliferation.

## INTRODUCTION

Stem cell-derived cardiomyocyte cell therapy has been established as a promising strategy for cardiac repair following myocardial infarction. Numerous groups have demonstrated both long-term engraftment and increased cardiac function in acute models of myocardial infarction in rodents and large animals ^1–7^. However, established techniques for transplanting human embryonic stem cell-derived cardiomyocytes (hESC-CMs) *in vivo* are hindered by small graft sizes, resulting from limited early cell retention and high rates of post-transplant cell death ^8–10^. Consequently, a high cell dose is required in order to achieve a therapeutic response after transplantation. Similar issues face other cell therapies, including those involving neural or islet cells ^11,12^. While tissue engineering strategies may address some of these limitations by implanting bulk tissues ^13–15^, other issues arise such as reduced electromechanical integration and the need for invasive implantation techniques ^5^. Thus, to facilitate the clinical translation and scalability of hESC-CM cell therapy, there is a need for methods to enhance graft size and to minimize the number of cardiomyocytes required for transplantation.

One strategy to address this is to enhance cardiomyocyte proliferation *in vivo* after transplantation. Notch signaling has been previously demonstrated to regulate cardiomyocyte proliferation ^16–22^, and full-length Notch ligands have been used to stimulate hESC-CM cell cycle activity *in vitro* ^19^. Direct cell-cell contact is typically required for Notch activation, as tension between the ligand and receptor likely exposes an extracellular cleavage site prior to releasing the Notch intracellular domain (NICD), which functions as a transcriptional cofactor. This presents a major hurdle in designing therapies utilizing Notch signaling, as ligands require immobilization and orientation through a signaling cell or surface to elicit a robust response ^23,24^. Previous studies have addressed this *in vitro* by activating Notch through ligand immobilization on plates or beads ^18,19,25,26^ or by utilizing viral overexpression systems ^17,18^, however these techniques are limited in their translational potential due to more complicated delivery techniques required ^27,28^.

An alternative approach that is compatible with cell-based therapy is to immobilize Notch ligands within an injectable biomaterial. Many injectable materials have been investigated for myocardial transplantation, including naturally occurring extracellular matrix (ECM)-derived proteins as well as synthetic biomaterials ^29,30^; however, few studies have modified the materials to immobilize signaling proteins in order to manipulate cell fate ^29,31^. Notch activation has been achieved in this context through a self-assembling peptide functionalized with a peptide mimic of the Notch ligand Jagged-1, however, these studies were limited to c-kit^+^ rat progenitor cells ^16^, now known to have minimal cardiogenic potential ^32,33^. We hypothesized that Notch ligand immobilization onto a natural, 3D scaffold would allow for transient activation of the Notch pathway in stem cell-derived cardiomyocytes, which could be used to promote proliferation and enhance engraftment after transplantation into a cardiac injury model. Thus, we sought to design an approach that would be compatible with established hESC-CM cell therapy techniques, using an injectable biomaterial that gels *in situ* to allow for needle delivery of hESC-CMs and the Notch ligand into the myocardial wall.

Here we have developed a novel approach to reduce the required therapeutic dose of cells for myocardial repair by promoting proliferation of injected cardiomyocytes via immobilized Notch signaling in a conveniently injectable hydrogel scaffold. We designed a collagen-based hydrogel with the immobilized Notch ligand Delta-1, which is used to promote the proliferation of engrafted cardiomyocytes after transplantation through activating the Notch signaling pathway. This Delta-1 functionalized hydrogel was first validated *in vitro*, where activating Notch signaling resulted in an increase in hESC-CM proliferation. The hydrogel was then used as an injectate for hESC-CM transplantation into rat myocardial infarcts, resulting in a 2-fold increase in engrafted cardiomyocyte proliferation and a corresponding 3-fold increase in graft size compared to controls. Furthermore, implantation of the Delta-1 functionalized hydrogel with a subtherapeutic dose of hESC-CMs led to maintenance of heart function at three months.

## RESULTS

### Delta-1 promotes Notch signaling and hESC-cardiomyocyte proliferation in 2D *in vitro*

Notch signaling activation with the Notch ligand Delta-1 was first confirmed in 2D using a U2OS CSLluc/ren luciferase reporter cell line (Fig. S1). The Delta-1 ligand was fused to an Fc domain, and this ligand was immobilized and presented in an oriented manner by binding it to anti-IgG, which had been adsorbed to the 2D tissue culture polystyrene (TCPS) plates. In this context, peak luciferase expression occurred between 48 and 72 hours with an increase of 3.5 ± 0.2-fold, 5.8 ± 0.2-fold, 6.1 ± 0.4-fold, and 4.6 ± 0.2-fold over uncoated TCPS and IgG control gel coated TCPS at 24, 48, 72, and 96 hours, respectively (Fig. S1A, p<0.005 at all timepoints) with expression returning to baseline by 7 days (Fig. 1A). This response was further enhanced with the addition of fibronectin (Fig. S1B), which was included to promote attachment of hESC-CMs in subsequent experiments. Culturing high purity hESC-CMs (>95% cTnT^+^) on Delta-1 coated surfaces (Delta) resulted in a significant increase in proliferation compared to hESC-CMs cultured on IgG control-coated surfaces (Control), measured by immunohistochemistry for double-labeled BrdU^+^/βMHC^+^ cells. This proliferative response was dose-dependent, with absolute proliferation rates for Delta increasing by 15.1 ± 3.1%, 22.0 ± 6.7%, and 11.5 ± 2.9% on 5 µg/ml, 10 µg/ml, and 20 µg/ml surfaces, respectively, versus Control (Fig. S1C). This enhanced cell cycle activity also corresponded with a significant increase in cell number, suggesting that the cardiomyocytes completed mitosis rather than becoming tetraploid. Addition of a gamma-secretase inhibitor, which inhibits cleavage and release of the NICD, blocked the increase in cell number induced by Delta, consistent with a Notch-dependent effect. Interestingly, proliferation rates in Control plates were also reduced by gamma-secretase inhibition, suggesting that Notch signaling is contributing to the basal proliferation rates of hESC-CM in monolayers (Fig. S1D).

**Figure 1.**
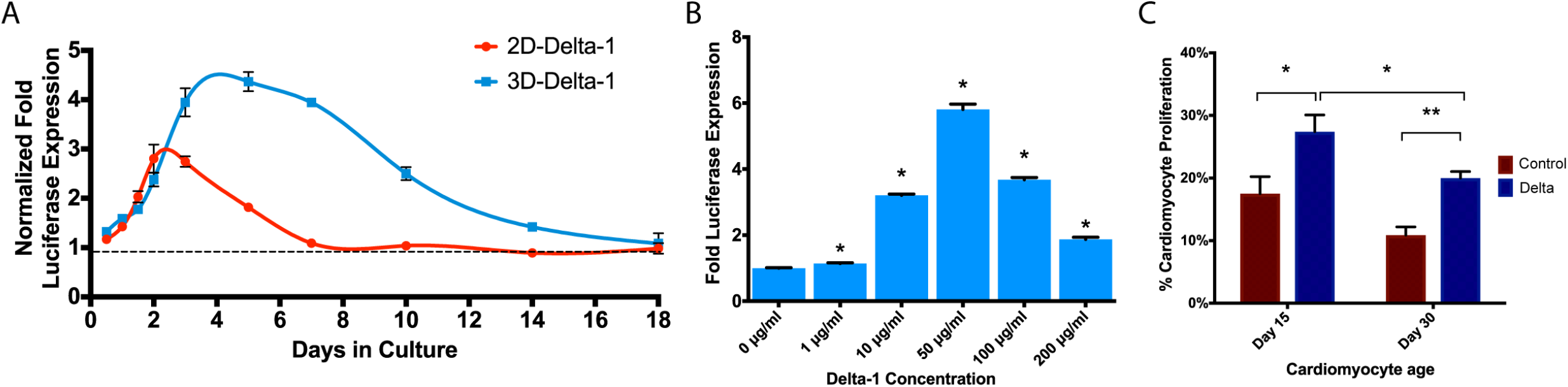
Prolonged Notch activation in 3D gels is dose-dependent and promotes hESC-CM proliferation. (A) Time course analysis of Notch-mediated luciferase expression in U2OS CSL luc/ren Notch reporter cell line indicates a prolonged Notch signal response in 3D culture conditions compared to the 2D Delta-1 platform. Luciferase signal is expressed normalized to IgG controls (2D or 3D) and plotted against days in culture. IgG controls were normalized to one (dashed line). (B) Dose-dependent Notch activation. Culturing U2OS CSL luc/ren Notch reporter cell line in the 3D Delta-1 gel results in Delta-1 dose-dependent activation of Notch signaling, as indicated by fold luciferase expression compared to IgG control gels. (C) HESC-cardiomyocytes in 3D engineered cardiac tissues proliferate in response to Delta-1 (Delta). Cardiomyocyte proliferation was measured by histology as double-positive βMHC^+^/BrdU^+^ cells, which results in a significant increase in response to Delta with day 15 cells and day 30 cells over IgG controls (Control). Note that, although proliferation slows over time, significant augmentation by Delta is still possible at day 30. For panel B, p-values were calculated using a one-way ANOVA with all samples compared to 0 µg/mL followed by Dunnett’s multiple comparisons test. For panel C, p-values were calculated using a multiple unpaired t test without assuming consistent SD. * indicates p<0.05, ** indicates p<0.005, and error bars denote SEM. See also Figure S1-2.

### Delta-1 promotes Notch signaling and hESC-cardiomyocyte proliferation in 3D engineered tissues *in vitro*

In order to achieve controlled Notch signaling in a 3D culture environment, we designed a collagen-based scaffold with immobilized Delta-1 by cross-linking anti-IgG to collagen and subsequently loading the gel with Delta-1 ligand (Fig. S2A). We then validated Notch activation *in vitro* by forming engineered tissues using either the U2OS CSLluc/ren reporter cells or hESC-CMs. While direct, unoriented conjugation of Delta-1 did not significantly increase Notch signaling over controls in 3D collagen gels, we found that binding Delta-1 through an intermediate anti-IgG protein allowed for ligand orientation and resulted in a 3.7 ± 0.2-fold increase over control gels (p<0.005), and a 3.1 ± 0.1-fold increase over unoriented Delta-1 (p<0.005) (Fig. S2B). This activation was further optimized by increasing ligand-collagen incubation time (Fig. S2C), which led to a significant and dose-dependent increase in Notch signaling compared to the established 2D ligand coating platform (Fig. 1A-B). Notch-driven luciferase expression in 3D Notch-gels peaked at day 5 with a 4.4 ± 0.2-fold increase over controls, and remained 2.5 ± 0.1-fold higher than 3D control gels at day 10 (Fig. 1A). Luciferase expression was still detectable after 2 weeks in 3D Delta-1 gels but declined back to baseline levels by day 18 (1.1 ± 0.2-fold). This represents a 3.9-fold increase in total Notch signal (area under curve) when compared to 2D platforms.

When hESC-CMs were seeded into the Delta-1 gels to form engineered cardiac tissues, we found hESC-CM proliferation was significantly increased compared to control (Fig. 1C). Tissues formed with hESC-CMs at day 15 after the initiation of directed differentiation showed a proliferative rate of 17.5 ± 2.7% in control tissues compared to 27.4 ± 2.7% in Delta-1 tissues (p=0.037). As expected, older hESC-CMs at day 30 after directed differentiation showed a lower basal rate of proliferation but also demonstrated a proliferative response to Delta-1 (20.0 ± 1.0% on Delta-1 compared to 10.9 ± 1.3% with control, p=0.0002). These data demonstrate that the 3D Delta-1 gel induces Notch signaling and increases proliferation in human cardiomyocytes in engineered heart tissues.

### Delta-1 gel increases graft size and vascularization in infarcted hearts

Following *in vitro* validation of the collagen gel with immobilized Delta-1, we next investigated whether Notch signaling could enhance proliferation and improve cellular engraftment *in vivo* in a model of myocardial infarction. We modified established protocols in our laboratory for transplanting hESC-CMs into the infarcted rat heart with pro-survival cocktail ^1,5,6^ to include gel functionalized with either IgG (Control) or Delta-1 (Delta), as 50:50 vol/vol of the injectate along with implantation of 10×10^6^ hESC-CMs. Unexpectedly, we found no significant difference in graft area (quantified from histology for βMHC) or infarct area (quantified from picrosirius red stain) identified between groups (Fig. S3A-B). A previous study suggests that Notch signaling is inhibited by cyclosporine A ^34^, one of the components of our pro-survival cocktail. Indeed, *in vitro* addition of pro-survival cocktail components, including cyclosporine, resulted in a 2-fold reduction in the Notch signaling response in Delta gels compared to standard media treatment (p<0.0001, Fig. S3C). To validate this *in vivo*, we excluded the pro-survival cocktail and performed a pilot study (n=2 per group) injecting a lower dose of cells (5 million cells per heart) in the same gel volume (either Control + hESC-CM or Delta + hESC-CM), and found a 2-fold enhancement in graft size in the Delta-gel (Figure S4). Based on these results, pro-survival cocktail was excluded from all subsequent studies.

In the 2-week pilot study, graft size in the Delta-1 group was comparable to control grafts (with pro-survival cocktail) at 1 month, despite having only half the cell dose and half the time for *in situ* proliferation. This suggests that when coupled with pro-proliferative Notch signaling, fewer implanted cells may be required to achieve myocardial remuscularization. To test this with greater rigor, we systematically assessed graft size at 1-month after transplantation using a lower cell dose. Five million hESC-CMs were transplanted into the infarcted rat myocardium in either Control or Delta gel (Control + hESC-CM and Delta + hESC-CM, respectively) (Fig. 2A, Fig. S5). Four weeks after implantation, human myocardial grafts were identified within the infarct regions by histology for βMHC (Fig. 2B-D). Implantation of Delta + hESC-CM resulted in a significant 3-fold increase in graft area compared to implantation of Control + hESC-CM, with grafts covering 3.0 ± 0.6% of the infarct region in Delta + hESC-CM and only 1.0 ± 0.2% in Control + hESC-CM (p=0.04, Fig. 2F). This corresponded to human myocardial grafts that comprised 1.0 ± 0.3% of the left ventricle in Delta + hESC-CM, a 3-fold increase over Control + hESC-CM (p=0.016, Fig. 2G). Consistent with the *in vitro* studies, Delta + hESC-CMs showed a higher level of cell cycle activity, identified by co-staining for BrdU and βMHC (animals were given a BrdU pulse at days 1, 4, 7, and 14 post-implantation). Delta + hESC-CM were 10.9 ± 0.5% double-positive for BrdU^+^/βMHC^+^ compared to only 4.8 ± 0.5% in Control + hESC-CM, corresponding to a 2.3-fold increase in proliferation (p=1.27E-05, Fig. 2H). Infarct area and anterior wall thickness was equivalent between groups (Fig. 3A-B). Vascularization was assessed by histological staining for CD31 to label endothelial cells, followed by quantifying microvessel density within the graft region. Delta + hESC-CM increased neovascularization by 4.4 ± 1.3-fold compared to Control + hESC-CM (p=0.04, Fig. 3G-H, J). We also observed a decreasing trend of macrophage content normalized to infarct area at 2 and 4 weeks; however due to high variability in the control group this was not statistically different (Fig. 3C-F, I).

**Figure 2.**
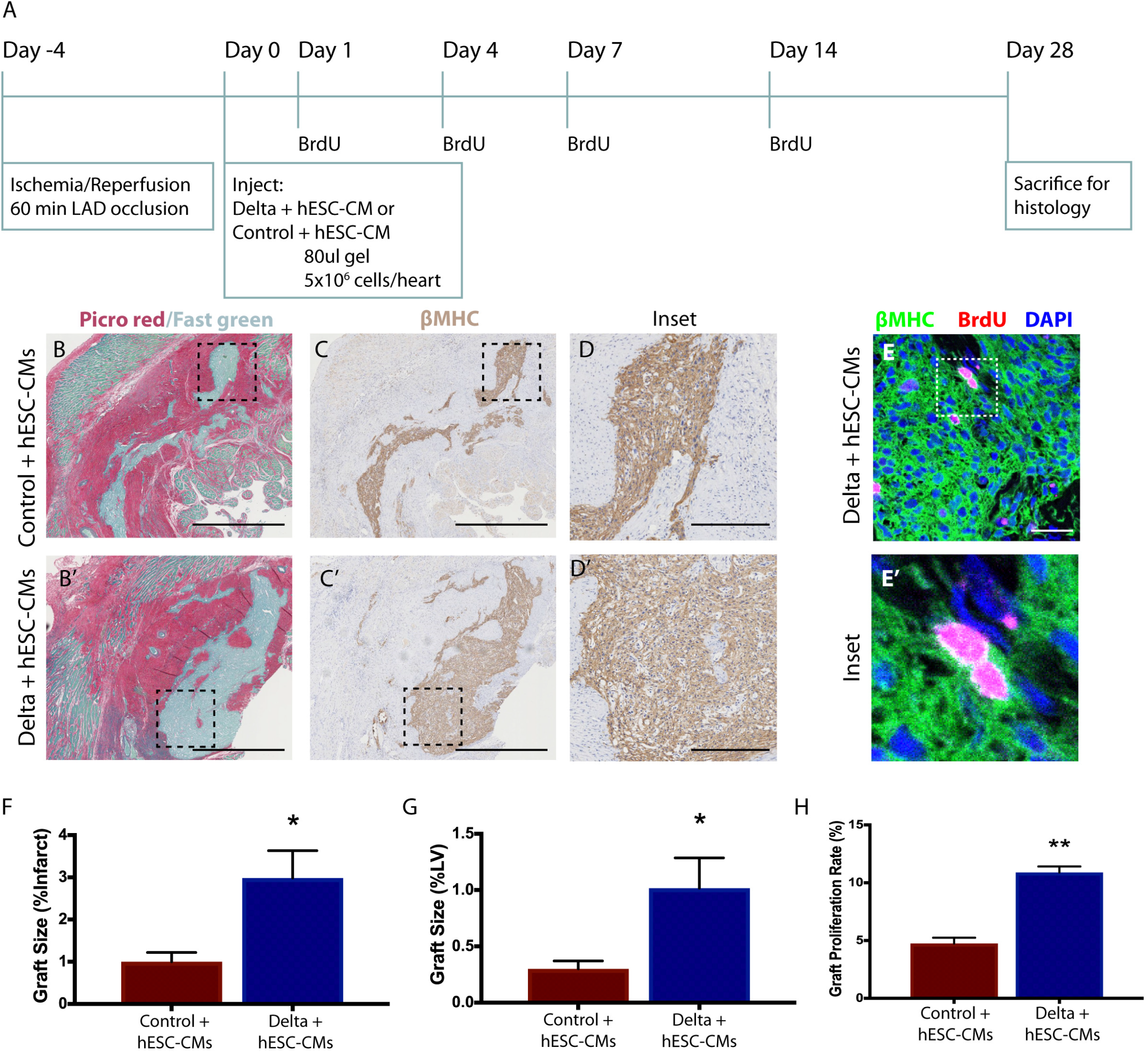
Notch signaling enhances hESC-CM engraftment and proliferation at one month post-implantation. (A) Experimental timeline. Four days after I/R injury, 5×10^6^ hESC-CMs were transplanted in either IgG or Delta-1 modified gel (Control + hESC-CMs and Delta + hESC-CMs, respectively), and tissues were harvested after 4 weeks. (B, B’) Collagenous scar area is identified by picrosirius red staining with a fast green counterstain to label healthy myocardium. Outline identifies region of interest. Scale bar = 1 mm. (C, C’) In serial sections, human myocardial grafts are identified by staining for beta myosin heavy chain (βMHC, brown) with hematoxylin counterstain. Scale bar = 1 mm. (D, D’) Regions of interest outlined in previous panels are shown at higher magnification with staining for βMHC. Scale bar = 250 µm. (E) Proliferation of transplanted hESC-cardiomyocytes is identified by histology. Tissue sections are stained with antibodies to detect beta myosin heavy chain (βMHC, green), BrdU (pink), and nuclear counterstain of DAPI (blue). Scale bare = 200 µm. Region of interest outlined in (E) is shown at higher magnification in (E’). (F) Graft area normalized to left ventricular area is significantly augmented in Delta + hESC-CMs. (G) Graft area normalized to infarct area is significantly augmented in Delta + hESC-CMs. (H) Engrafted cardiomyocyte proliferation identified by βMHC^+^/BrdU^+^ cells is significantly enhanced in Delta + hESC-CMs. For panels F-H, P-values were calculated using an unpaired two-tailed t test and * indicates p<0.05, ** indicates p<0.0001, and error bars represent SEM. See also Figure S3-5.

**Figure 3.**
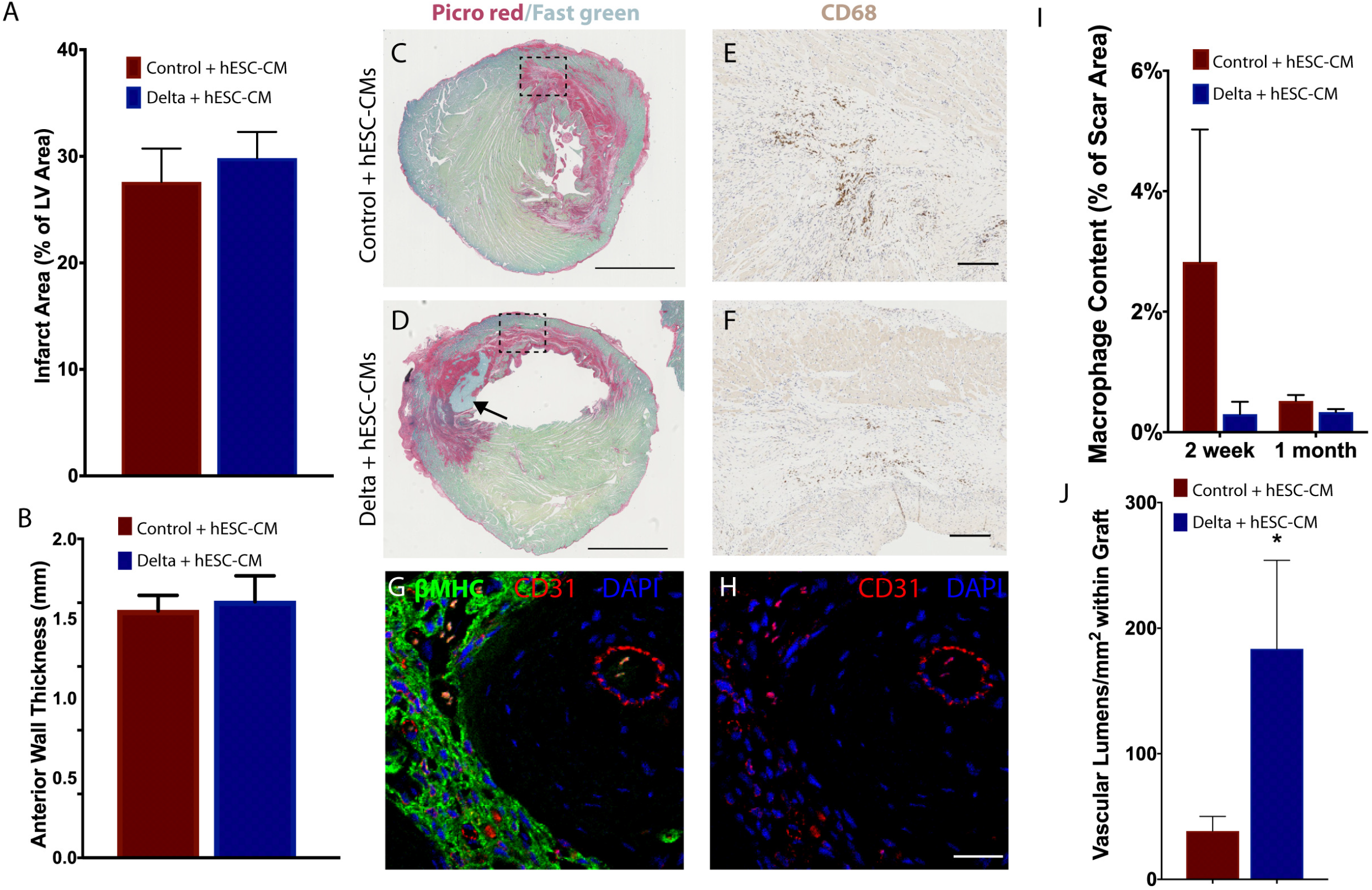
Notch signaling enhances vascularization with similar infarct size at one month post-implantation. (A) Average infarct area is assessed by histology and normalized to left ventricular (LV) area at 4 weeks. (B) Anterior wall thickness by histology is shown for an area approximately 4 mm from the apex of the heart. Values are in mm. (C, D) Picrosirius red and fast green counterstain was used to identify fibrotic regions that are quantified in (A). Representative images are shown for both Control + hESC-CMs (C) and Delta + hESC-CMs (D), where the black arrow identifies a region of human myocardial graft within the infarct. Scale bar = 2.5 mm. (E, F) Inflammatory response at 4 weeks by CD68 staining normalized to infarct area. Outlined regions of interest in C, D are shown at higher magnification in E, F. Serial sections are stained with CD68 antibody to label monocytes and macrophages and visualized with DAB (brown). Scale bar = 200 µm. (G, H) Host-derived vessels are identified by staining with CD31 antibody (red) with a double stain for βMHC (green) to identify hESC-cardiomyocyte grafts within the infarct regions. Scale bare = 200 µm. (I) Level of inflammatory response is expressed as CD68^+^ area normalized to scar area by picrosirius red at 2 weeks and 4 weeks. Values for 2 week time point were obtained through pilot study experiments shown in Fig. S4. (J) Level of neovascularization is quantified as CD31^+^ lumens within βMHC^+^ graft regions. P-values were calculated using an unpaired two-tailed t test. For panels A-B, F-G, * indicates p<0.05 and error bars indicate SEM.

### Transplantation with Delta-1 gel improves cardiac function in a low cell-dose model

Previous studies have established that transplantation of 10 million hESC-CMs were required to achieve therapeutic functional benefit in the infarcted athymic rat heart ^6^. Due to increased graft proliferation at one month, implantation of only 5 million hESC-CMs in the Delta-1 functionalized hydrogel resulted in increased graft size despite a lower cell dose. We hypothesized that this subtherapeutic cell dose could be lowered further and still result in therapeutic benefit post-infarct.

To provide a stringent test of the ability of the Delta-1 functionalized hydrogel to improve cardiac repair in the long-term with hESC-CMs, we delivered a subtherapeutic dose of cells (2.5 million cells per heart) to the infarcted myocardium and followed the animals by serial echocardiography for three months. Two additional gel-only control groups were added to evaluate the possible effects of the gel without the presence of cells, resulting in four groups total: Control gel–no cells, Delta gel–no cells, Control gel + hESC-CMs, and Delta gel + hESC-CMs (Fig. 4A). Consistent with the data at one month post-transplant, implantation of Delta + hESC-CMs resulted in a significant increase in βMHC^+^ graft area compared to implantation of Control + hESC-CMs (Figs. 4B-C). The graft size was smaller in both groups compared to our previous studies, likely due to the reduced cell dose. As anticipated based on our results at the one month endpoint, Delta + hESC-CMs exhibited a higher cell cycle activity, as identified by staining for double-positive BrdU^+^/βMHC^+^ cells after a series of four BrdU pulses over the first two weeks post-implantation (Fig. 4D).

**Figure 4.**
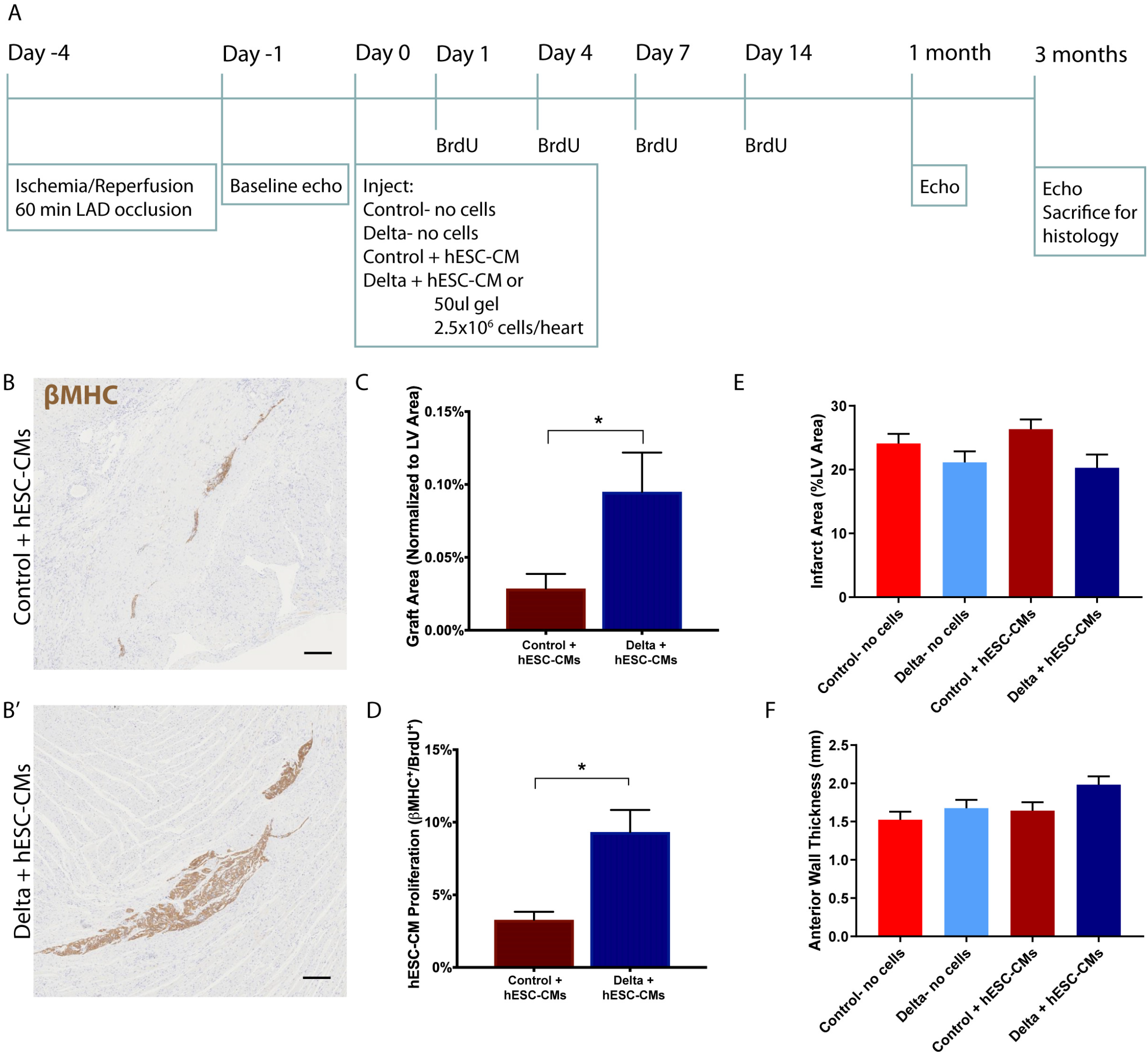
Human myocardial graft area is significantly increased three months after implantation with Notch signaling. (A) Experimental timeline. Four days after ischemia/reperfusion injury, 2.5×10^6^ hESC-CMs were transplanted within the IgG control gel or the Delta-1-gel (Control + hESC-CMs and Delta + hESC-CMs, respectively), with two additional gel-only control groups (Control-no cells and Delta-no cells). (B, B’) Representative images of hESC-CM grafts identified by staining for beta myosin heavy chain (βMHC, brown) with hematoxylin counterstain for Control + hESC-CM (B) and Delta + hESC-CM (B’). Scale bar = 200 µm. (C) βMHC^+^ graft area is normalized to LV area. There is a substantial increase in graft area with the Delta + hESC-CMs. (D) Proliferating hESC-CMs are identified by double-labeled βMHC^+^/BrdU^+^ cells, and quantification is shown here. There is a significant increase in Delta + hESC-CM proliferation. (E) Infarct area, quantified by picrosiriurs red area, is normalized to LV area. There is a modest, non-significant reduction in infarct size with Delta + hESC-CM treatment. (F) Anterior wall thickness is shown in mm. There is a modest, non-significant increase in anterior wall thickness in the Delta + hESC-CMs. For panels C-F, * indicates p<0.05 and error bars indicate SEM. P-values were calculated for panels C-D using an unpaired t-test and for E-F using a one-way ANOVA followed by Sidak’s multiple comparisons test comparing Control-no cells vs. Delta-no cells, Control-no cells vs. Control + hESC-CMs, Delta-no cells vs. Delta + hESC-CMs, and Control + hESC-CMs vs. Delta + hESC-CMs.

Three months after implantation, there was no significant difference in histological infarct size (Fig. 4E). Infarcts occupied 20.3 ± 2.1% of the LV in Delta + hESC-CMs hearts and 21.1 ± 1.7% in Delta– no cells hearts, compared to 26.3 ± 1.5% and 24.1 ± 1.5% in the Control + hESC-CMs and the Control– no cells groups (Fig. 4E). There was no difference in anterior wall thickness in the Delta + hESC-CMs compared to all other groups (Fig. 4F).

To determine whether the increased graft area observed in the Delta + hESC-CMs group impacted global heart function, echocardiography was performed on all animals at baseline post-injury, and at one and three months post-implantation. Measurements were conducted by two independent, blinded observers per animal and time point and showed high interobserver agreement (Fig. S7A-B). All groups had comparable left ventricular end-diastolic dimension and fractional shortening at the post-infarct baseline echocardiogram, suggesting comparable baseline infarct sizes prior to cell and gel implantation (Fig. 5A, Fig. S6B, D). In hearts receiving Control gel + hESC-CMs, there was a significant reduction in fractional shortening from baseline to one month (from 29.4 ± 1.0% to 26.2 ± 1.1%), and further reduction to 24.4 ± 0.7% at three months (compared to baseline, p=0.039 at one month, p=0.0002 at three months, Fig. 5B). This is similar to the progression of heart failure in Control gel—no cells group (Fig. 5B) and indicates that the dose of cells in this group is indeed subtherapeutic. In contrast, animals in the Delta gel + hESC-CMs group showed no progression of heart failure after transplantation (28.1 ± 0.9% FS at baseline, versus 28.7 ± 1.2% and 29.3 ± 1.0% at one and three months, respectively, Fig. 5B). The change in fractional shortening from baseline trended towards improvement at one month (Fig. 5C) and was significantly improved at three months when animals received Delta + hESC-CMs compared to Control gel – no cells and Control gel + hESC-CMs (Fig. 5D). The Delta gel—no cells group also showed no significant decline in function over time with a mean fractional shortening of 28.3 ± 1.0%, 28.0 ± 0.9%, and 26.2 ± 0.8% at baseline, one month, and three months, respectively (Fig. 5B). This suggests a benefit of Notch signaling on host cells in the infarcted heart, but unlike the Delta gel + hESC-CM group, there was no statistically between-group improvement in function in the Delta gel—no cells group, indicating that hESC-CM provide additional benefit. Left ventricular end-diastolic and end-systolic dimensions increased across baseline and one and three months in all groups (Fig. S6A, C), with no significant between-group differences (Fig. S6B, D). Taken together, these data indicate that the Delta-1 gel augments the ability of an otherwise subtherapeutic dose of human cardiomyocytes to improve systolic function of the infarcted heart, in association with enhanced graft size and improved graft vascularization.

**Figure 5.**
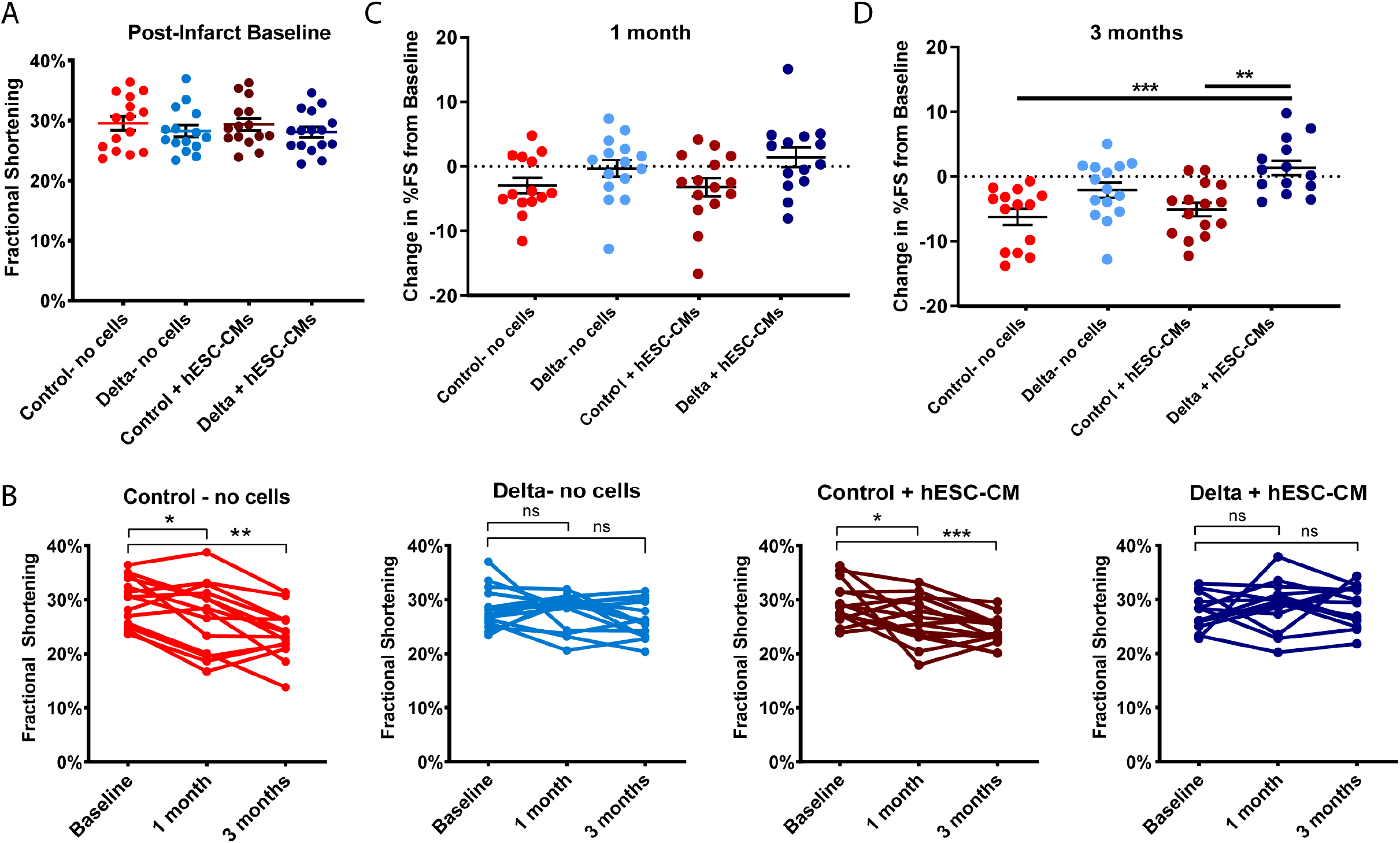
Notch signaling maintains heart function at three months with implantation of a subtherapeutic dose of hESC-CMs. (A) Echocardiography was performed at baseline after ischemia/reperfusion injury with no differences between groups, indicating a similar extent of injury prior to implantation. (B) Individual fractional shortening data from baseline to one and three months are shown for all four implant groups. Note the decline in fractional shortening over time in Control-no cells and Control+ hESC-CMs, whereas fractional shortening is maintained in Delta-no cells and Delta + hESC-CMs. Paired t-tests were performed comparing one and three months with baseline within each group. Change in percent fractional shortening from baseline, at one month (C) and three months (D). Each data point represents the change in a heart’s function at the corresponding time point, with group means and standard error shown for each group. Control – no cells and Control + hESC-CMs show a significantly greater decrease in fractional shortening at three months compared to the Delta + hESC-CMs as determined by a one-way ANOVA followed by Sidak’s multiple comparisons test comparing Control-no cells vs. Delta-no cells, Control-no cells vs. Control + hESC-CMs, Control – no cells vs. Delta + hESC-CMs, Delta-no cells vs. Delta + hESC-CMs, and Control + hESC-CMs vs. Delta + hESC-CMs. For all panels, ns = p>0.05, * p<0.05, ** p<0.005, *** p<0.0005 and error bars represent SEM. See also Figure S6-7.

## DISCUSSION

Cell therapy methods for cardiac repair are hindered by small graft sizes resulting from limited retention and high cell death after implantation. This has traditionally required transplanting a high cell dose of hESC-CMs into the myocardium, which is a hurdle for translation of this therapeutic approach to the clinic. To address this, we developed a method to increase the proliferation of hESC-CMs after transplantation via Notch signaling activation, which resulted in increased graft size and maintained heart function with a normally subtherapeutic cell dose.

Notch ligand immobilization in 2D has been used to successfully mediate cell fate decisions in hESCs and hESC-CMs ^19^, however we found that similar 2D surfaces begin to lose signal activity within 2-3 days in culture. In contrast, we demonstrated in Fig. 1 that immobilizing a Notch ligand onto a 3D scaffold allowed for significant Notch signaling activation for up to 2.5 weeks of *in vitro* culture. This data indicates a robust increase in total Notch signal gained from the 3D system compared to 2D, however it also indicates that Notch signaling activation decreases back to baseline levels over time. This is important particularly for *in vivo* translation, because overly proliferative cardiomyocytes have been shown to contribute to decreased global heart function when the proliferative stimuli are overexpressed long-term in the heart ^35,36^. This highlights the importance of the transient proliferative cues achieved here with the Delta-1 gel, which we believe allows for a balance between cardiomyocyte proliferation and the eventual maturation of *de novo* cardiomyocytes needed for functional improvement.

Interestingly, by implanting hESC-CMs with the Delta-1 gel we could achieve human myocardial grafts that, two weeks after implantation, were comparable to the graft sizes we’ve generated in previous studies that used twice the cell input number at one month ^5,6^. To determine the ability of our Delta-1 gel to improve heart function with an otherwise subtherapeutic dose of cells, we reduced the cell dose for the three month functional experiments, where only 2.5×10^6^ hESC-CMs were implanted instead of 5×10^6^ hESC-CMs implanted in the one month experiment and the 10×10^6^ hESC-CMs implanted in comparable published studies ^5,6^. Importantly, the increase in hESC-CM proliferation and subsequent increase in graft size was identified at both one and three months post-transplantation of hESC-CMs in the Delta-1 gel, suggesting graft stability at later time points even with a reduced cell input. Indeed, Delta + hESC-CMs showed a maintenance of heart function over the three month time period following implantation, where Control + hESC-CMs showed functional decline (Fig. 5). Additionally, Delta + hESC-CMs showed statistically significant improvement in the change in fractional shortening from baseline when compared with Control + hESC-CMs. This data confirms the injection of a subtherapeutic dose of cells and the ability of our Notch signaling gel to promote proliferation of grafted cells and prevent functional decline.

Based on results from previous studies by our group where animals received cell-free injections after an infarction ^6^, we anticipated that the Control gel—no cells group would show a decline in heart function over one month and continue to decline over three months, which is consistent with the data shown here (Fig. 5). Due to the previously reported evidence of Notch-mediated effects on cardiac repair in mice and zebrafish ^20,37–41^, we also hypothesized that the transient Notch signal in the Delta groups may stimulate an endogenous repair response in the rat host myocardium, resulting in some benefit gained by the Delta-1 signal without the presence of hESC-CMs in the Delta– no cells group. While Delta-no cells did not show a functional decline over time (Fig. 5B) and there was a trend for improved change in fractional shortening from baseline compared to the Control gel—no cells animals, this was not statistically different. Interestingly, the largest increase in fractional shortening from baseline at one and three months was identified in rats receiving Delta + hESC-CMs, suggesting a combinatorial impact of the Notch activation and cell therapy in this context.

In conclusion, we developed a Notch-signaling hydrogel to promote the proliferation of hESC-CMs, which was validated *in vitro* using tissue engineering and translated *in vivo* as an injectate for hESC-CMs to increase graft size and maintain heart function. Implanting hESC-CMs in the Delta-1 gel resulted in a 2-fold increase in the proliferation of engrafted cardiomyocytes and corresponded to a 3-fold increase in graft size compared to cell transplantation with a control gel at one month, and importantly, prevented functional decline with a subtherapeutic dose of cells at three months. Collectively, these results demonstrate the use of a Delta-1 functionalized hydrogel as a therapeutic approach to enhance hESC-CM graft size after implantation into injured heart through transiently stimulating proliferation, allowing an otherwise subtherapeutic dose of hESC-CMs to improve systolic function after an acute myocardial infarction.

## MATERIALS AND METHODS

### 2D Tissue Culture Platforms with Immobilized Notch Ligands

To achieve oriented immobilization of recombinant human Delta-1-Fc ^24^, tissue culture polystyrene culture plates were pre-coated with 20 mg/ml anti-IgG (human, Fc-specific, Sigma) for 1 hour at 37°C. Wells were washed with PBS, blocked with 20% FBS in PBS for 1 hour at room temperature, and subsequently incubated overnight with the Notch ligand Delta-1 (courtesy of the Bernstein Lab, FHCRC) or human IgG (Sigma) to achieve oriented immobilization. A similar method has been previously described ^23^. 2D cell culture experiments used Delta-1 ligand or IgG control at a 10 µg/ml unless otherwise noted. Prior to cell culture, wells were washed three times with PBS. In 2D experiments using hESC-CMs, 5 µg/ml fibronectin (Invitrogen) was added during overnight ligand incubation to promote cardiomyocyte attachment to the tissue culture surfaces.

### Collagen Modification and Notch Ligand Immobilization

Carbodiimide chemistry was used to immobilize Delta-1/Fc onto solubilized collagen similar to previously described methods ^42,43^, where free carboxyl groups on the collagen are reacted to free amine groups to bind a secondary protein; anti-IgG in all experiments, except for un-oriented ligand immobilization experiments that used either IgG or Delta-1 bound directly to collagen. To activate the collagen, a stock concentration of 15 mg/ml rat tail collagen type 1 was dissolved in 0.1% acetic acid, diluted to 2.5 mg/ml using RPMI-1640 cell culture medium, and reacted with 1-ethyl-3-(dimethylaminopropyl) carbodiimide hydrochloride (EDC)/*N*-sulfohydroxysuccinimide (sulfo-NHS) on ice for 1 hour, with a 10-fold molar excess of EDC (ThermoFisher) to free carboxyl groups and 5 mM Sulfo-NHS (ThermoFisher). The solution was mixed periodically during the incubation period using a chilled 1cc syringe. After 1 hour, 2-mercaptoethanol (2-ME) was added at 20 mM to inactive the EDC, and the pH was adjusted to 7.0 – 7.2 using 1M NaOH. Next, 200 μg/ml of anti-human IgG (Fc domain-specific) was mixed with the collagen and incubated for 48 hours at 4°C. Following the 48 hour incubation, 100 µg/ml human recombinant Delta-1 or human IgG (Sigma) was added, and the solution was incubated for another 24 hours at 4°C. Prior to the addition of cells, the solution was brought to a final concentration of 1.5 mg/ml collagen using cell culture medium (DMEM for U2OS luciferase characterization studies or RPMI-1640 for cardiomyocyte experiments), 1x HEPES, 30 vol/vol% unmodified rat tail collagen 1 (4 mg/ml stock, ThermoFisher), 1x Matrigel, and the pH was adjusted to 7.4 using 1M NaOH. This entire process occurred with reagents, pipettes, and syringes maintained on ice.

### Luciferase-based Analysis of Notch Signaling

CSL-luciferase U2OS osteosarcoma cells (courtesy of Dr. Randall Moon’s Lab, University of Washington) and the Dual-Luciferase Reporter Assay System (Promega) were used for 2D and 3D Notch signaling activation experiments *in vitro*. U2OS cells were engineered to express constitutively active renilla and firefly luciferase under control of CSL (CBF1, Su(H), and Lag-1) expression (cells are referred to as U2OS CSLluc/ren). U2OS CSLluc/ren cells were cultured in uncoated 10 cm TCPS plates and fed every 2-3 days with DMEM supplemented with 10% FBS, 2 mM l-glutamine, and 100 units/ml penicillin/ 100 µg/ml streptomycin, and passaged at 80% confluence using 0.25% trypsin-EDTA.

For 2D experiments, U2OS CSLluc/ren were replated in triplicate onto IgG- or Delta-1-coated wells in a 96-well plate at a density of 5k cells/well in standard culture medium. For 3D gel experiments, 25k U2OS CSLluc/ren were resuspended in 30 µl of IgG- or Delta-modified collagen and allowed to gel at 37°C in 96-well plates (in 3-6 replicates) for 1 hour before adding an additional 200 µl of cell culture medium. Bioactivity of implanted gels was assessed using gels formed *in vitro* with CSLluc/ren reporter cells and harvested at 72 hours for luciferase analysis. For luciferase analysis, medium was removed and 100 or 200 µl of 1x Passive Lysis Buffer (Promega) was added to 2D and 3D wells, respectively. Cells were lysed during a 10 min incubation followed by trituration and 2-3 repetitive freeze/thaw cycles at −80°C. 3D gels required an additional trituration step to fully disrupt the gels, which was accomplished using a handheld tissue homogenizer inserted into each sample tube and triturated for 15 sec. Luminescence was recorded following the Dual-Luciferase Reporter Cells protocol (Promega). Briefly, 20 µl per sample was loaded in triplicate into optical 96-well plates, and background fluorescence was recorded over 3 individual reads on a plate luminometer. 20 µl of Luciferase Assay Reagent II (LARII) was added to each well and firefly luciferase luminescence was recorded as before, followed by the addition of 20 µl Stop and Glo Reagent to each well and luminescence recording. Notch-driven luciferase data is represented as normalized firefly to renilla luminescence for each sample, expressed as the fold expression over the corresponding IgG control (normalized to 1).

### hESC-Derived Cardiomyocyte Culture and Differentiation

Undifferentiated RUES2 hESCs were maintained as previously described ^5^ on Matrigel™ (Corning) in mouse embryonic fibroblast (MEF)-conditioned media supplemented with 5 ng/ml basic fibroblast growth factor (bFGF) (Peprotech). Cardiomyocyte differentiation was performed as described previously using a combination of small molecules and growth factors ^5^. A high-density cell monolayer was pre-treated with 1 µM CHIR99021 for 6-24 hrs (Cayman Chemical), and induced with 100 ng/ml Activin A (R & D Systems) and 1x Matrigel in RPMI-1640 with B27 Supplement minus insulin (Life Technologies). Medium was changed after 18 hours to RPMI-1640 with B27 Supplement (minus insulin) supplemented with 1 µM CHIR99021 and 5 ng/ml BMP4 (R & D Systems). After two days media was changed and supplemented with XAV939 (Tocris) for an additional 48 hours. After day 7 of differentiation, RPMI-1640 with B27 supplement containing insulin (Life Technologies) was used to maintain cells. Beating was typically observed between days 7 and 10, and medium was changed every 2-3 days thereafter. Unless otherwise noted, cardiomyocytes were cryopreserved on day 21-24 of differentiation for long-term storage and thawed 2-3 days prior to implantation to allow for recovery ^44^. For cryopreservation, cells were heat shocked for 1 hour at 42**°**C the day prior. After a 1 hour pretreatment with 10 µM ROCK inhibitor Y-27632 (Tocris), cardiomyocytes were harvested by a brief incubation with EDTA and dispersed into single cells using 0.25% trypsin-EDTA (Life Technologies). Cells were washed and resuspended in CryoStor (Sigma), added to cryovials, and frozen to −80**°**C in a controlled rate freezer with a decrease in temperature of 1**°**C/minute before transfer to liquid nitrogen for long-term storage. To thaw, cryovials were agitated briefly at 37**°**C, collected in RPMI-1640 supplemented with 200 U/ml DNAse (VWR), and washed with basal medium. Cardiomyocytes were replated onto Matrigel-coated culture dishes in RPMI-1640 with B27 containing insulin and supplemented with 10 µM Y-27632 for the first 24 hrs. Flow cytometry for cardiac troponin T (1:100, Thermo Scientific) was used to characterize cardiomyocyte population purity after cardiac differentiation and again at the time of harvest for implantation.

### *In vitro* Cardiomyocyte Proliferation Experiments

HESC-CMs were replated into wells of a 4-well Nunc^TM^ chamber slide at 60k cells/well (Thermo Scientific) following immobilization of either IgG or Delta-1 and supplemented with 5 µg/ml fibronectin as described above. Culture media was changed the following day including 10 µM BrdU (Sigma), and cells were fixed 24 hours later with 4% paraformaldehyde. For experiments where cell number was the primary endpoint, 24-well plates were used instead of chamber slides, and were coated with Delta-1 or IgG as described above. For these experiments, high-purity cardiomyocytes resulting from the differentiation protocol described above (97.1 ± 0.9% cTnT^+^ by flow cytometry) were replated at a density of 200k cells/well, medium was changed every other day, with or without the addition of the gamma-secretase inhibitor DAPT (Sigma) at 5 µM, and cells were counted on a hemocytometer using Trypan Blue dye at day 7 post-replating. Engineered cardiac constructs were formed using methods previously described by our group ^45–47^. HESC-CMs were removed from monolayer culture using 0.25% trypsin-EDTA, washed twice with serum-free medium, resuspended in the modified collagen with IgG (control) or Delta-1, and added into sterile PDMS molds using a 1cc syringe for 30 µl gels with 0.5×10^6^ cells/gel in a PDMS mold. Collagen gels were allowed to polymerize by incubation at 37°C for 1 hour, followed by the addition of RPMI-1640+B27 containing insulin supplemented with 10 µM Y-27632 for the first 24 hrs. Medium was changed every other day, with 10 µM BrdU included 24 hours prior to fixation with 4% paraformaldehyde.

### Pro-survival Cocktail Experiments

Initial implantation studies which included, including pro-survival cocktail were performed as previously described ^5,6^ using a 1:1 mix of pro-survival cocktail with the modified collagen for implantation. As per the pro-survival cocktail protocol, animals received daily subcutaneous injections of 5 mg/kg cyclosporine A for 7 days, starting the day prior to implantation (for experiments shown in Fig. S3A-B). For the set of experiments shown in Fig. S3C, we investigated the effect of pro-survival cocktail on Notch signaling *in vitro* using the U2OS luciferase assay described above. U2OS cells were resuspended in DMEM containing the pro-survival cocktail which includes 100 µM ZVAD (benzyloxycarbonyl-Val-Ala-Asp(O-methyl)-fluoromethyl ketone), 50 nM Bcl-X_L_ BH4, 200 nM cyclosporine A, 100 ng/mL IGF-1, and 50 µM pinacidil. In these studies and in the implantation studies including the pro-survival cocktail, the 50% vol/vol Matrigel described in the original reference ^5,6^ was replaced with the modified collagen gel (immobilized with IgG or Delta-1). To mimic the daily cyclosporine injections given to rats during the first week following implantation, daily medium was supplemented with 200 nM cyclosporine A.

Because we found the pro-survival cocktail to have a negative impact on Notch signaling, it was omitted from all other implantation experiments and was is only included in the experiments shown in Fig. S3.

### Surgical and Implantation Procedures

All animal procedures were performed in accordance with the US NIH Policy on Humane Care and Use of Laboratory Animals and were approved by the UW Institutional Animal Care and Use Committee. Male Sprague Dawley athymic rats (8-12 weeks old, 250 – 300 g) (Envigo) were weighed and anesthetized with an intraperitoneal injection of 68.2 mg/kg ketamine (Zoetis) and 4.4 mg/kg xylazine (Akorn Animal Health), intubated, and mechanically ventilated. Core body temperature was maintained at 37°C by placing the rat on a warming pad and monitoring rectal temperature every 15 min. A local block of 1 mg/kg lidocaine (Hospira) and 1 mg/kg bupivacaine (Hospira) was injected subcutaneously at the incision site prior to surgery start. A thoracotomy was performed to expose the heart, and ischemia was induced by occluding the left descending coronary artery with a 7-0 suture for 60 min followed by reperfusion. The chest was closed aseptically and animal recovery was monitored. An analgesic dose of 1 mg/kg sustained-release buprenorphine (Zoopharm) was given subcutaneously before rehousing the animals the same evening. All animals received post-operative care twice-daily for two days following surgery and were monitored daily until the second surgery.

Four days after ischemia/reperfusion (I/R), rats were again weighed and given 1 mg/kg sustained-release buprenorphine subcutaneously at least 1 hour prior to surgery start. Animals were then anesthetized with 2.5-5% isofluorane, mechanically ventilated, and a second thoracotomy was performed to expose the heart. For one month studies (Fig. 2-3), 5×10^6^ freshly-harvested (not cryopreserved) RUES2 hESC-CMs were delivered in 80 µl volume of either the IgG-collagen (Control + hESC-CMs) or Delta-1-collagen gel (Delta + hESC-CMs) in 2 injections within the infarct region using a 26g needle. For pilot subtherapeutic cell dose 2 week study (Fig. S4), a normally subtherapeutic dose of 5×10^6^ freshly-harvested (not cryopreserved) RUES2 hESC-cardiomyocytes was delivered in 80 µl volume of Notch signaling or control gel, (Delta + hESC-CMs and Control + hESC-CMs, respectively). For three month studies (Fig. 4-5 and S6-7), 2.5×10^6^ (freshly-harvested, not cryopreserved) RUES2 hESC-cardiomyocytes were delivered in 50 µl volume of Notch signaling or control gel, (Delta + hESC-CMs and Control + hESC-CMs, respectively) and additional control groups received either cell-free Delta-1 gel (Delta-no cells) or cell-free IgG control gel (Control-no cells). After cell injection, the chest was closed aseptically and animal recovery was monitored. All animals received post-operative care twice-daily for 2 days followed by monitoring 3-4 times per week until their endpoint. BrdU was administered via intraperitoneal injection at 10 mg/kg on days 1, 4, 7, and 14 post-cell injection. Echocardiography, using either a GE Vivid 7 Dimension Ultrasound or Visual Sonics Vevo 2100, was performed during the three month studies at 4 days after infarction (baseline) and at one and three months post-cell injection. To perform echocardiography, animals were lightly anesthetized with 1-2.5% isoflurane, and the left ventricular (LV) end diastolic dimension (LVEDd), LV end systolic dimension (LVEDd), fractional shortening (%FS) and heart rate were recorded. Animals were euthanized at their endpoint using a lethal overdose of Beuthanasia (Merck Animal Health) administered via intraperitoneal injection. Hearts were collected, washed in cold PBS, perfused with PBS to flush out blood, and soaked in 150 mM KCl. During this time, a small sample of intestine was collected from each animal to serve as a BrdU-positive control.

### Immunohistochemical Analysis

After euthanasia, hearts were fixed in 4% paraformaldehyde overnight, sectioned, processed, and stained with the appropriate primary and secondary antibodies. Infarcted myocardium was visualized using picrosirius red with a fast green counterstain, and picrosirius red area was normalized to left ventricular area during analysis. Human cardiomyocyte grafts were identified by staining for beta myosin heavy chain (βMHC) (hybridoma supernatant, ATCC #CRL-2046) and visualized using either Alexa Fluor-488 goat anti-mouse (1:100, Molecular Probes) or an avidin-biotin goat anti-mouse antibody (1:100, Vector Labs) developed with diaminobenzidene (Vector Labs). To analyze BrdU incorporation, slides were first stained with the βMHC primary antibody as described above and then treated with 1.5 N HCl, followed by overnight incubation with a peroxidase-conjugated anti-BrdU primary antibody (1:40, Roche). Slides were then incubated with Alexa Fluor-488 goat anti-mouse (1:100, Molecular Probes) for βMHC and AF594-tyramide (Thermo Fisher) to amplify BrdU. Cardiomyocyte proliferation was assessed by counting double-labeled BrdU^+^/βMHC^+^ cells, which was analyzed on images obtained on a Nikon A1R confocal microscope. Vasculature was identified using CD31/PECAM-1 primary antibody (1:100, Novus) and macrophages/monocytes were visualized using CD68 primary antibody (1:100, Serotec).

### Statistical Measurements

All histological measurements were performed using ImageJ and statistical analysis were performed using Excel or Prism Graphpad. Statistical tests performed and significance values are indicated in figure legends for each experiment. All values are reported as means, and error bars represent standard error of the mean (SEM).

## ACKNOWLEDGEMENTS

The authors thank Dr. Irwin Bernstein for providing the Delta-1 ligand and for helpful discussions regarding Notch signaling, Daniel Burnham and Alexander Moon for assistance with immunohistochemistry, Gabriel Rush for assistance with data collection, and Dr. Lil Pabon and Dr. Hans Reinecke for insightful scientific discussions. We gratefully acknowledge funding from the Foundation Leducq Transatlantic Network of Excellence and NIH grants R01HL146868, R01HL128362, U54DK107979, and P01HL094374 (to CEM). KAG was supported by NIH training grant T32EB001650 and an NSF Graduate Research Fellowship. We gratefully acknowledge the Tom and Sue Ellison Stem Cell Core of the Institute for Stem Cell and Regenerative Medicine for use of cell culture space and equipment. This research was supported by the Cell Analysis Facility Flow Cytometry and Imaging Core in the Department of Immunology at the University of Washington. We would like to acknowledge the Mike and Lynn Garvey Cell Imaging Lab at the Institute for Stem Cell and Regenerative Medicine (UW) and its director Dale Hailey for assistance with sample imaging and analysis. Histopathology work was provided by the Pathology Research Service laboratory at the University of Washington. We would like to thank Megan Larmore and Brian Johnson in the University of Washington Histology and Imaging Core for their assistance with whole slide digital scanning.

## AUTHOR CONTRIBUTIONS

KAG designed and performed experiments, analyzed data, and wrote the manuscript; KAM performed experiments, analyzed data, and edited the manuscript; AM performed surgeries, echocardiography, and edited the manuscript; XG analyzed data and edited the manuscript; CEM supervised all aspects of the study, designed experiments, edited the manuscript, and obtained funding for the research.

