## Supplementary Information for "Delta-1 functionalized hydrogel promotes hESC-cardiomyocyte graft proliferation and maintains heart function post-injury"

**Contents:**

**Supplementary Figure 1:** Immobilized Delta-1 promotes Notch-mediated luciferase expression in 2D.
**Supplementary Figure 2:** Optimization of 3D gel protocol.
**Supplementary Figure 3:** Addition of pro-survival cocktail inhibits proliferation promoted by Notch signaling gel.
**Supplementary Figure 4:** Pilot study for reducing required cell dose.
**Supplementary Figure 5:** Gel and cell characterization prior to implantation.
**Supplementary Figure 6:** Left ventricular internal dimensions do not vary among groups by echocardiography.
**Supplementary Figure 7:** Echocardiography interobserver variability.

**Figure S1. Related to Figure 1.** Immobilized Delta-1 promotes Notch-mediated luciferase expression in 2D. (A) After culturing U2OS CSL luc/ren Notch reporter cell line on the 2D platform, Notch-driven luciferase expression (Delta) significantly increases relative to tissue culture polystyrene surfaces (Untreated) and IgG treated (Control) controls, peaking between 48 and 72 hrs. (B) Addition of fibronectin (FN) during ligand immobilization in 2D does not negatively impact Notch response, and in fact results in a slight increase in Notch signaling by CSLluc/ren luciferase analysis (Delta). (C) HESC-cardiomyocytes proliferate in response to immobilized, oriented Delta-1 (Delta) or IgG (Control) in 2D with inclusion of FN. There is a dose-dependent effect on cardiomyocyte proliferation, measured by histology as double-positive βMHC$^{+}$/BrdU$^{+}$ cells, with diminishing proliferation at the highest ligand density. (D) Human cardiomyocyte number increases after culture on 2D Delta-1 surfaces (Delta) compared with IgG coated surfaces (Control). The Notch inhibitor (gamma-secretase inhibitor, GSI) blocks Delta-induced proliferation and also reduces baseline proliferation. For all panels, * indicates p<0.05 and error bars denote SEM.

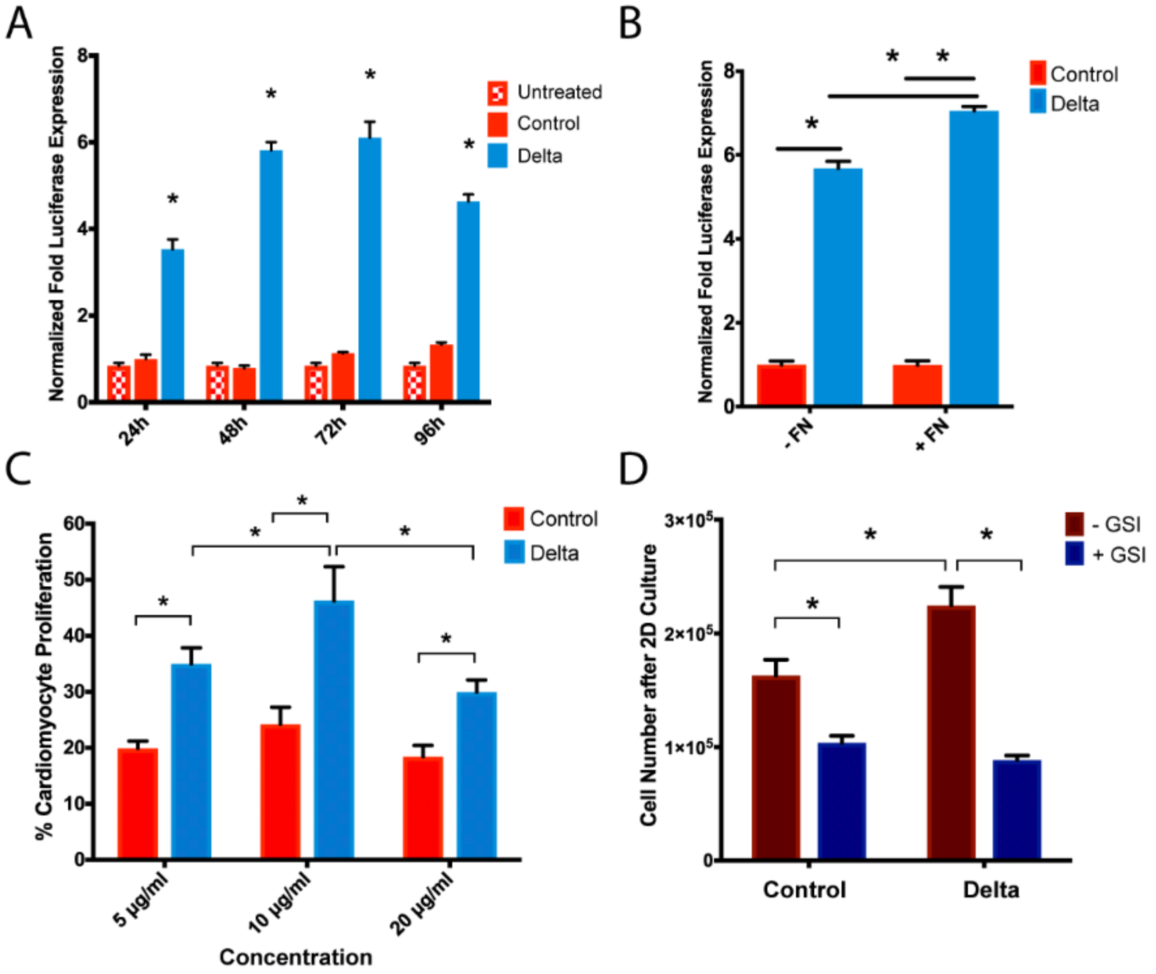

**Figure S2. Related to Figure 1.** Optimization of 3D gel protocol (A) 3D immobilization of Delta-1 or IgG as a control using 1-ethyl-3-(dimethylaminopropyl) carbodiimide hydrochloride (EDC)/*N*-sulfo-hydroxysuccinimide (sulfo-NHS) chemical cross-linking. After the stable collagen-anti-IgG conjugate is formed, the solution is incubated with Delta-1 (Delta) or IgG (Control) for ligand immobilization and orientation. (B) Delta-1 ligand orientation via anti-IgG binding is required to achieve detectable Notch-mediated luciferase expression in 3D gels. Samples were analyzed after 48 hrs in culture. (C) Increasing the duration of anti-IgG binding time facilitates higher Delta-1 immobilization as measured through Notch-driven luciferase expression. Samples were analyzed after 48 hrs in culture. For all panels, * indicates $p<0.05$ and error bars denote SEM.

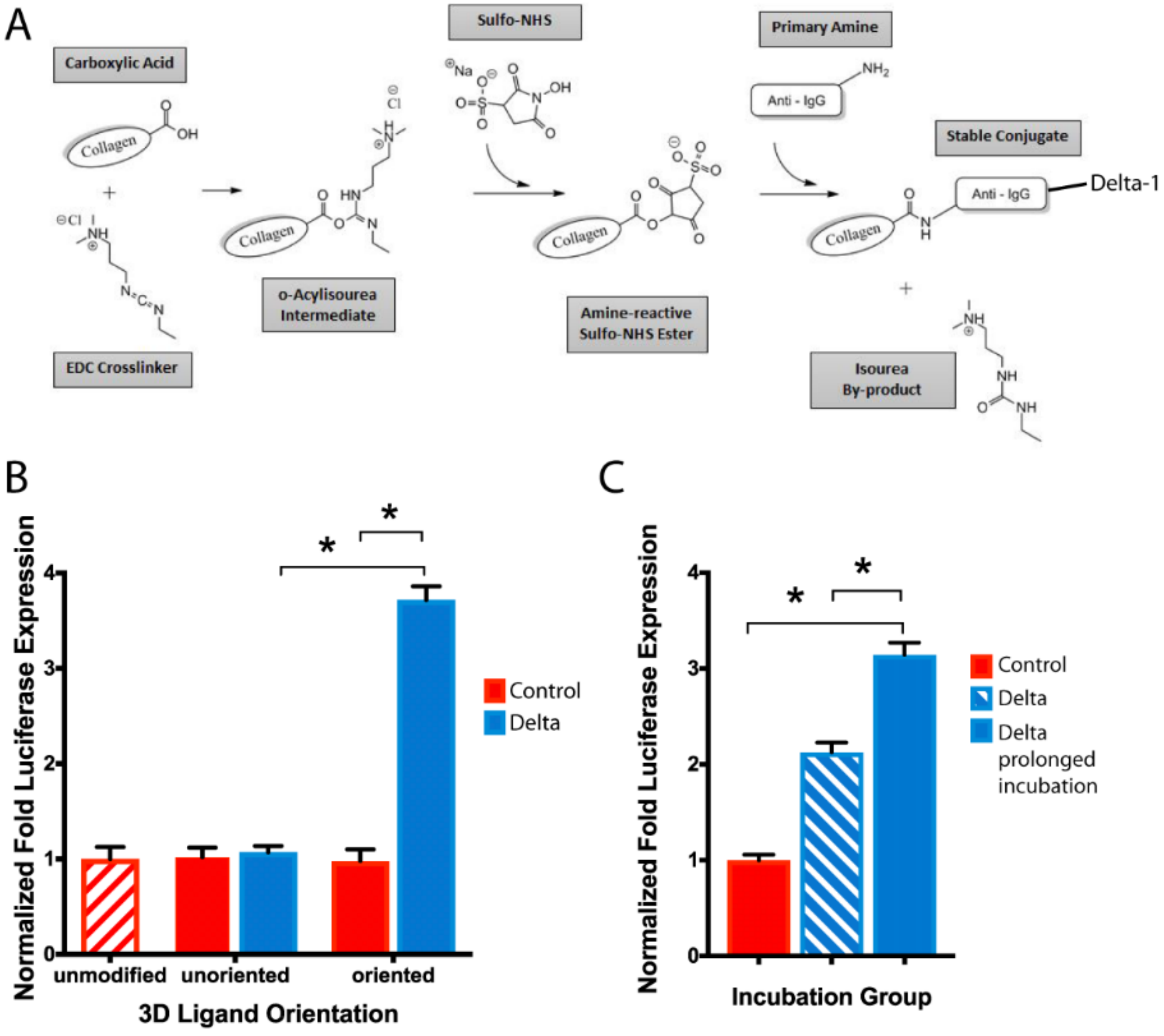

**Figure S3. Related to Figures 2, S2.** Addition of pro-survival cocktail inhibits proliferation promoted by Notch signaling gel. (A) Initial one-month study, where pro-survival cocktail was used and $10x10^6$ hESC-CMs were implanted. Grafts were identified by staining for βMHC and quantified with normalization to infarct area. Each data point indicates the total graft area within one heart, normalized to infarct area with horizontal lines indicating the group average. There was no statistical difference between groups. (B) Infarct area analysis shows no significant differences in scar size by histology (measuring picrosirius red infarct area, normalized to left ventricular area) at 4 weeks. (C) Pro-survival cocktail inhibits Notch signaling by luciferase analysis *in vitro*. Engineered tissues were formed using the U2OS CSL luc/ren Notch reporter cell line and modified collagen gels with either Delta-1 or IgG control. Tissues were treated with pro-survival cocktail (PSC) or standard media (control). For all panels, * indicates $p<0.05$ and error bars denote SEM.

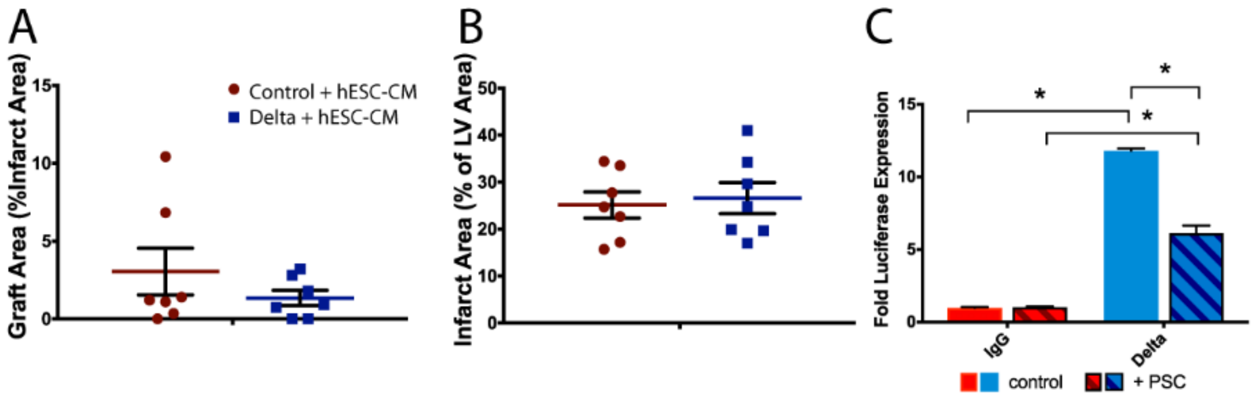

**Figure S4. Related to Figures 2, 3, S2.** Pilot study for reducing required cell dose. (A) Timeline for 2 week pilot study where animals received 5x10$^6$ hESC-CMs + IgG (Control + hESC-CMs) or Delta (Delta + hESC-CMs) gel. (B) Sum total of βMHC$^+$ graft area normalized to LV area across all analyzed tissue sections. Because n=2, no error bars are shown. (C) βMHC$^+$ graft area is normalized to fibrotic tissue area (picrosirius red area). Because n=2, no error bars are shown. (D) Fibrosis is identified in the myocardium by picrosirius red for collagen and healthy myocardium is counterstained with fast green. Scale bar = 5mm. (E) Transplanted human cardiomyocytes are identified with βMHC and visualized with DAB (brown) with a hematoxylin counterstain to quantify graft area. Scale bar = 100 μm.

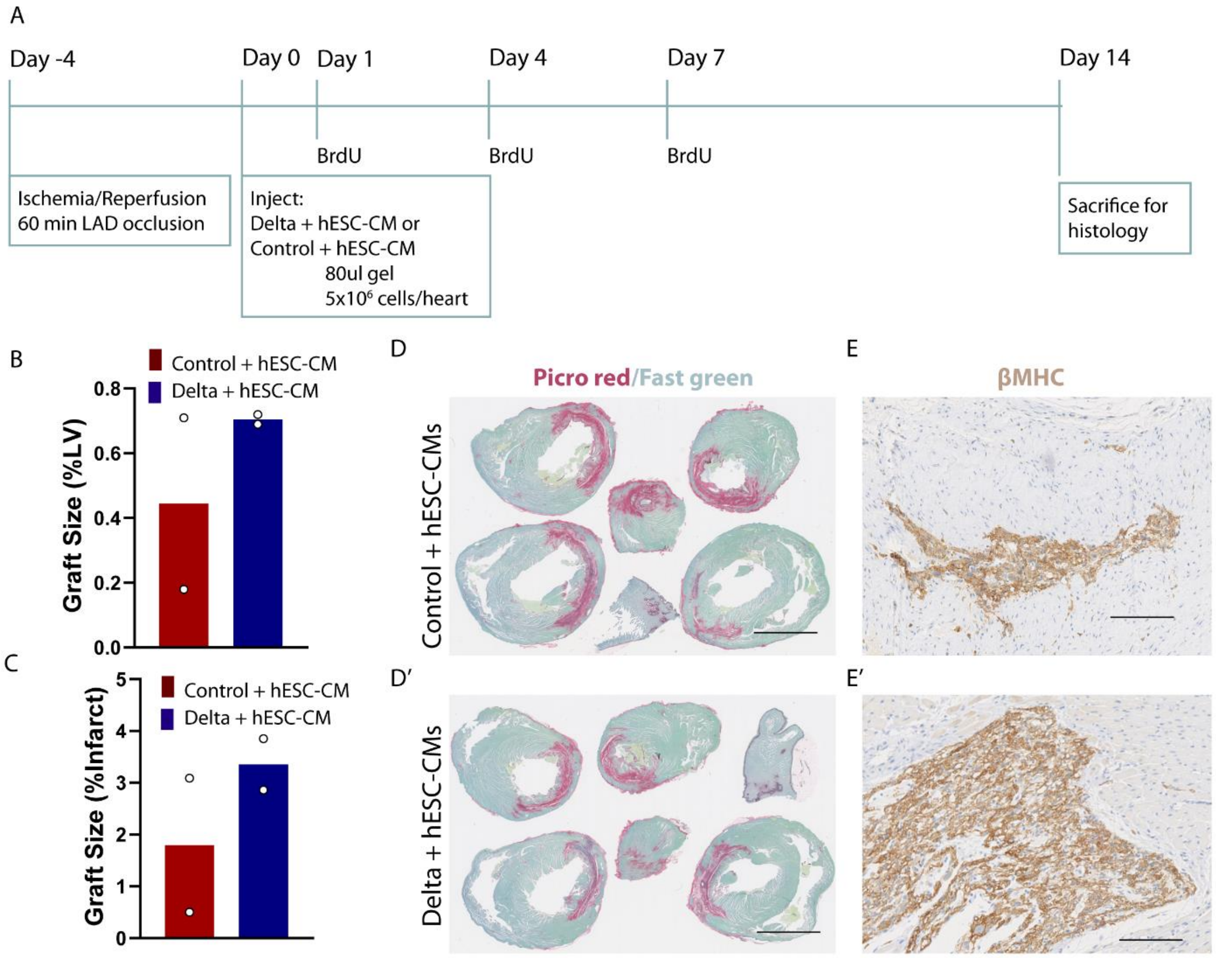

**Figure S5. Related to Figures 2, 3.** Gel and cell characterization prior to implantation. (A) Implanted gels bound with Delta-1 (Delta) produced a Notch-mediated increase in luciferase expression. Data is shown as fold increase of firefly vs. renilla luciferase, and normalized to IgG controls (Control). *indicates p<0.05, and error bars represent SEM. (B) Implanted hESC-cardiomyocytes were greater than 70% cTnT+ by flow cytometry. The cell population Forward Scatter (FSC-A) and Side Scatter (SSC-A) plots are shown on the left. Cardiac troponin T (cTnT) is detected by PE, where data is shown as a histogram of PE expression. The isotype control (red) is used as a negative gating control for the stained population (blue).

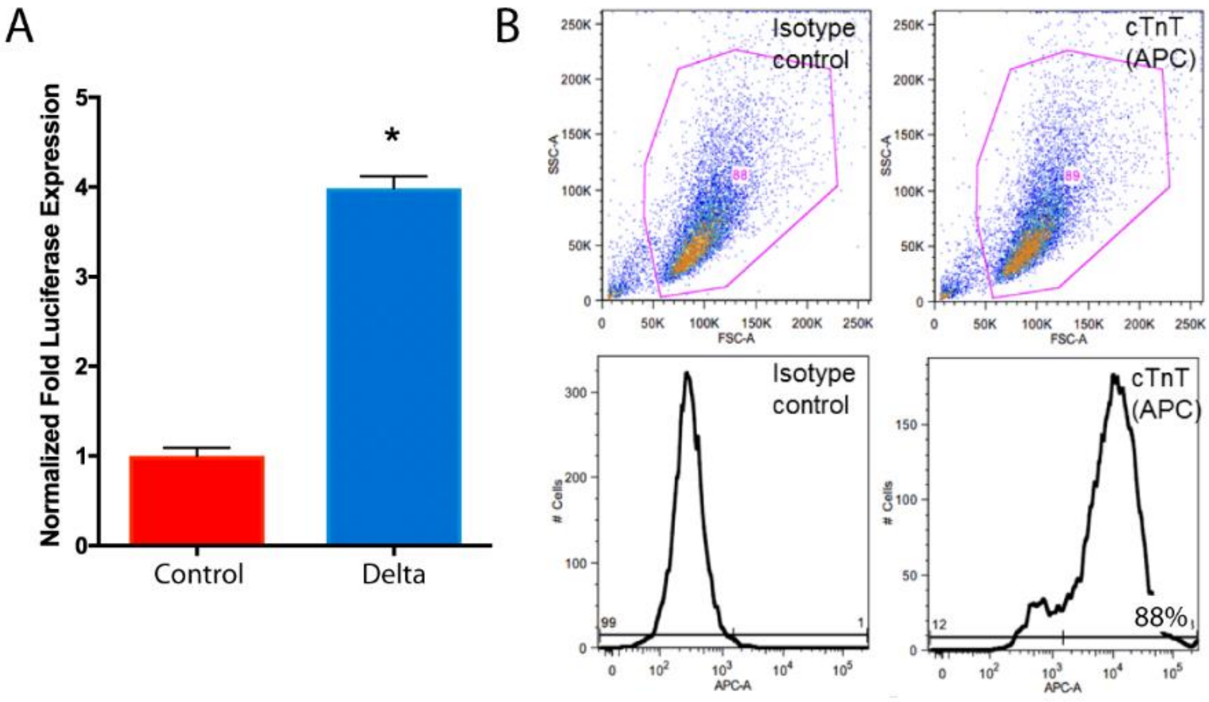

**Figure S6. Related to Figure 5.** Left ventricular internal dimensions do not vary among groups by echocardiography. Individual traces of left ventricular internal dimensions (LVID) are shown at baseline, one, and three months for each group at diastole (LVIDd, A) with group averages summarized in (B). Left ventricular internal dimensions are shown at systole (LVIDs) for individual hearts (C) with group averages shown in (D). For all panels, ns = p>0.05, * p<0.05, ** p<0.005, ***p<0.0005, ****p<0.0001 and error bars represent SEM. P-values were calculated by 2-way ANOVA followed by Tukey's multiple comparisons test.

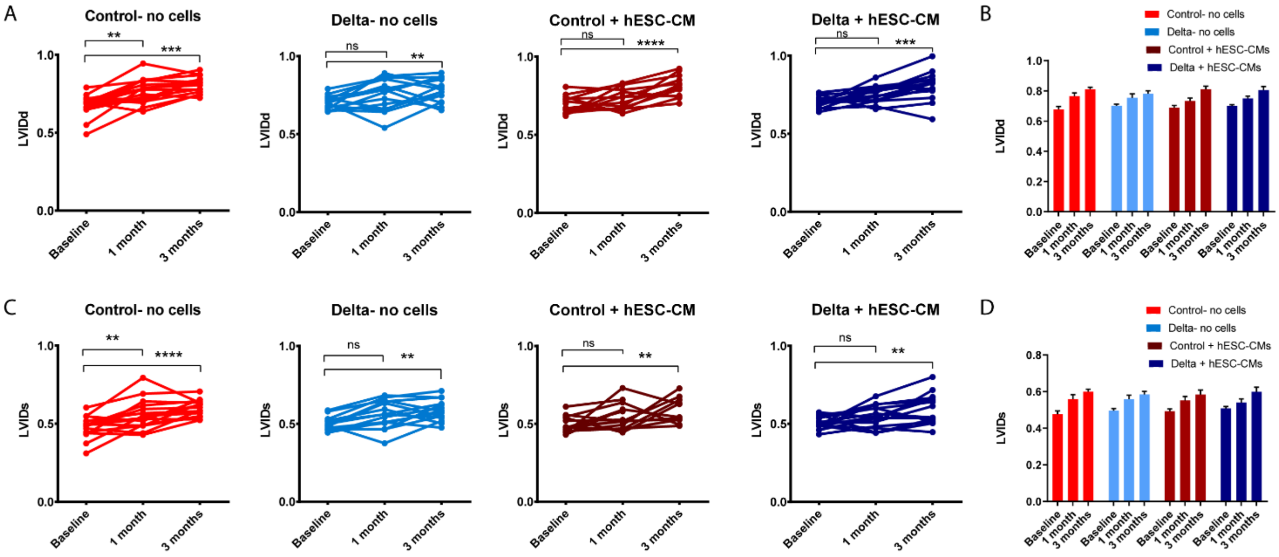

**Figure S7. Related to Figure 5.** Echocardiography interobserver variability. (A) Two independent, blinded scientists performed analysis of echocardiography with the correlation of measurements shown here for all time points (n=135). The best fit line equation and $R^2$ value are shown on the graph with Pearson correlation coefficient shown below. Surgery and echo analysis for 14 animals from the Control- no cell and Delta- no cell groups was performed at a later time than the initial experiment and on a different ultrasound. The correlation of observer measurements is shown separately for these animals at all time points (n=42) in (B).

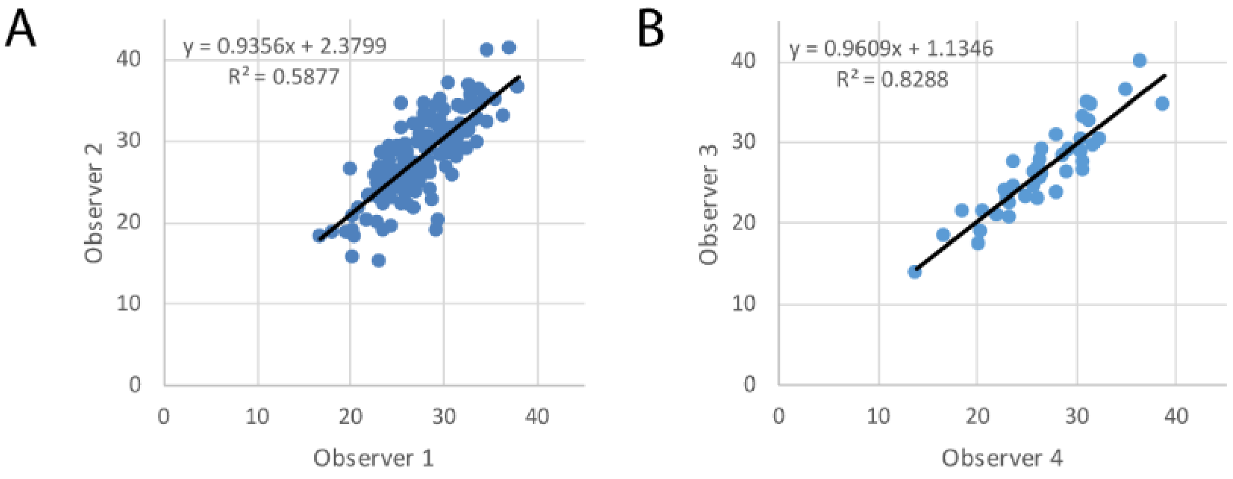


Pearson correlation coefficient 0.767

Pearson correlation coefficient 0.910
